# Distinct and Shared Impacts of Virulence Plasmids on the Phenotype and Transcriptome in Convergent Carbapenem-Resistant and Hypervirulent *Klebsiella pneumoniae*

**DOI:** 10.1101/2025.11.10.687661

**Authors:** Mingju Hao, Xiaodi Cui, Liya Feng, Ke Liu, Xiaohong Shi, Tengfei Long, Sarah E. Rowe, Yi-Tsung Lin, Liang Chen

## Abstract

The global rise of convergent carbapenem-resistant and hypervirulent *Klebsiella pneumoniae* (CR-hvKp) represents a major clinical challenge, yet the role of virulence plasmids (pVirs) in shaping bacterial physiology and pathogenicity remains incompletely understood. Using a CRISPR-Cas9–based curing system, we precisely eliminated pVirs from two clinical CR-hvKp strains with distinct genetic backgrounds, ST23-KL1 (*bla*_NDM-1_) and ST11-KL64 (*bla*_KPC-2_). Loss of pVir conferred fitness advantages in vitro, reduced capsule production and hypermucoviscosity, and promoted biofilm formation, while markedly attenuating virulence in murine sepsis models. Despite this reduction, the pVir-cured ST23-KL1 strain retained higher virulence than the pVir-cured ST11-KL64 strain, underscoring the contribution of chromosomal background to pathogenic potential. Transcriptomic profiling revealed both shared and strain-specific transcriptional responses to pVir deletion, with broader perturbations observed in the ST11-KL64 strain. pVir removal had limited effects on antibiotic MICs. Complementation experiments further demonstrated differential regulatory roles of the *rmpADC* and *rmpA2D2* operons in capsule expression and hypermucoviscosity across the two strains. Together, these findings establish pVirs as central determinants of CR-hvKp virulence and highlight complex host–plasmid interactions that influence bacterial adaptation and pathogenicity.

**IMPORTANCE:** The emergence of carbapenem-resistant and hypervirulent *Klebsiella pneumoniae* (CR-hvKp) poses a critical threat to global health, yet the contribution of virulence plasmids (pVirs) to bacterial fitness and pathogenicity remains poorly defined. By employing a CRISPR-Cas9–based curing strategy, we dissected the role of pVirs in two genetically distinct CR-hvKp strains and uncovered their multifaceted impact on capsule production, hypermucoviscosity, biofilm formation, and virulence. Our findings reveal that pVir loss confers fitness advantages in vitro while attenuating virulence in vivo, with strain-specific transcriptional responses and differential regulation by *rmp* operons. These results underscore the complex interplay between plasmid-encoded and chromosomal determinants in shaping CR-hvKp pathogenicity and adaptation, offering mechanistic insights that may inform future therapeutic strategies targeting plasmid-mediated virulence.

## INTRODUCTION

Hypervirulent *Klebsiella pneumoniae* (hvKp) is a highly pathogenic microorganism with the ability to cause infections in both community and hospital settings (1, 2). First recognized as a cause of community acquired pyogenic liver abscesses in Asia in the 1980s (3), it has now spread globally and causes a variety of infections, including liver abscess, bacteremia, pneumonia and soft tissue infections (4, 5). Classic *K. pneumoniae* (cKp) strains, in contrast to hvKp, have long been recognized as opportunistic pathogens, primarily associated with hospital-acquired infections in immunocompromised patients. The cKp strains are often multidrug-resistant (MDR), with resistance to multiple antibiotic classes, including beta-lactams, fluoroquinolones, and aminoglycosides. The emergence of carbapenem-resistant *K. pneumoniae* (CRKP) in classic Kp strains has further exacerbated the challenge of treating these infections, making them a critical concern in healthcare settings worldwide (6).

Historically, hvKp strains were susceptible to most antibiotics but now convergent multidrug-resistant and hypervirulent *K. pneumoniae* (MDR-hvKp), including those are resistant to carbapenems, are emerging, posing serious health concerns worldwide (7). The emergence of MDR-hvKp mainly occur via two major mechanisms: (i) acquisition of resistance genes by hvKp strains, such as the acquisition of carbapenemase gene in hvKp ST23-KL1 clone; (ii) acquisition of virulence plasmid by classic multidrug-resistant strains, such as the KPC producing ST11-KL64 clone (8–13).

The hypervirulent phenotype of hvKp has largely been attributed to the presence of a pK2044-like virulence plasmid (abbreviated as pVir in this study), which harbors the regulator of mucoid phenotype ADC genes (*rmpADC* and *rmpA2*), aerobactin (*iut*) and salmochelin (*iro*) siderophore biosynthetic genes and their cognate receptor genes (4, 14, 15). pVir-mediated hypercapsule formation, hypermucoviscosity (HMV), and hypersiderophore production are major virulence determinants of hvKp. In the hvKp K2 reference strain KPPR1, *rmpA* functions as an autoregulatory factor, activating the expression of downstream *rmpC* to stimulate capsule production and *rmpD* to drive the HMV phenotype (16–18). RmpD then binds to Wzc, a tyrosine kinase involved in capsule biosynthesis, which is essential for capsule polymerization and export. This interaction leads to the production of more consistent, longer polysaccharide chains, thereby leading to the HMV phenotype (18, 19). RmpA2, which shares approximately 80% identity with RmpA, plays a similar role as *rmpA* in enhancing capsule production (20, 21). In addition, a novel *rmpD2* gene was found downstream of *rmpA2* in most pVir plasmids, which contributes to HMV in the ST11-KL64 strain (22). The siderophore associated genes include aerobactin synthesis operon *iucABCD*, salmochelin production gene cluster *iroBCDN*, and the outer membrane ferric aerobactin receptor gene *iutA* (23). Siderophore increases the ability of hvKp to acquire iron from the host (1), of which aerobactin is the dominant siderophore produced by hvKp (24). The virulence plasmid also harbors gene *peg-344*, which encodes a putative transporter that is required for the full virulence of some hvKp in site-specific infection (2, 25). In addition, the pVir-borne tellurite resistance operon (*ter*) has been found to contribute to the fitness of gut colonization (26).

Despite being a hallmark of hvKp strains, the phenotypic and transcriptional impacts of pVir on different *K. pneumoniae* hosts remain unclear, partly due to the difficulty of generating isogenic pVir-cured strains in diverse clinical *K. pneumoniae* backgrounds. To address this gap, we employed a CRIPR-Cas9 mediated pCasCure plasmid-curing strategy to generate isogenic pVir-cured mutants. We then characterized the resulting phenotypic alterations and transcriptomic responses. Our findings reveal both shared and strain specific impacts of pVir in two distinct convergent carbapenem-resistant hvKp (CR-hvKp) backgrounds, providing new insights into the complex interplay between pVirs and host genomes in the evolution and pathogenicity of CR-hvKp strains.

## RESULTS

### Bacterial strains and case description

Two representative CR-hvKp isolates with distinct genetic backgrounds were selected for this study: JNQH373, an ST23 KL1 hvKp strain carrying the *bla*_NDM-1_ carbapenemase gene, and JNQH97, an ST11 KL64 CR-hvKp strain harboring the *bla*_KPC-2_ carbapenemase gene (Table 1). JNQH373 (ST23-KL1) represents a canonical hvKp lineage that has acquired a carbapenem resistance gene, whereas JNQH97 (ST11-KL64) exemplifies an emerging convergent MDR strain that has gained a pVir.

**Table 1.**
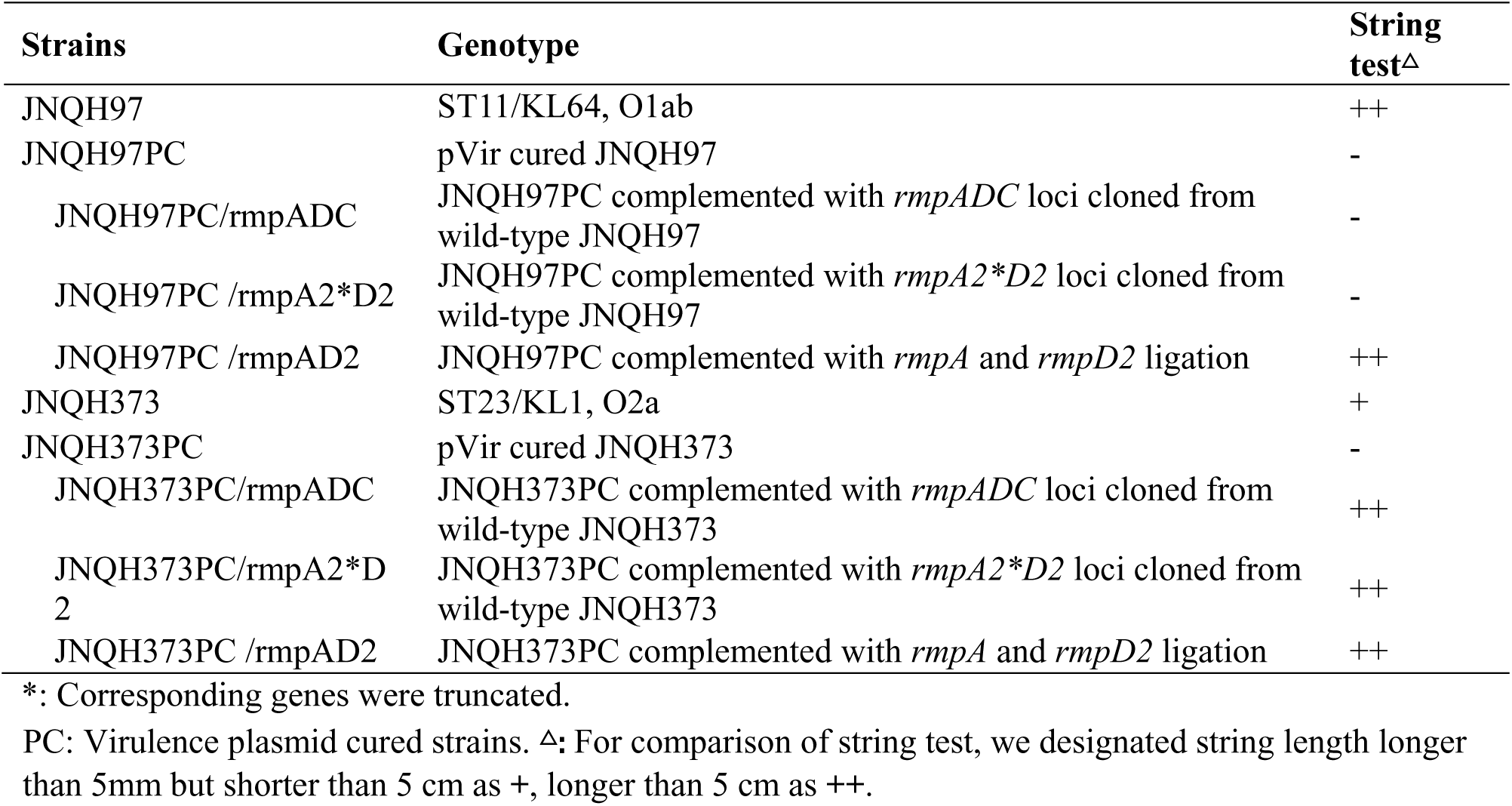
Strains used in this work.

Strain JNQH97 was recovered from a sputum sample of a female patient in her late fifties who was admitted to the intensive care unit (ICU) of a tertiary hospital in Eastern China in August 2014. She had a medical history of colon cancer, high blood pressure, and diabetes mellitus. Two days after admission, JNQH97 was isolated from a sputum sample, and three days after admission, she developed sepsis. During her seven-day hospitalization, she received multiple antimicrobial agents, including piperacillin-tazobactam and meropenem as empirical therapy. Despite treatment, her condition continued to deteriorate. The patient ultimately chose to discontinue medical care and was discharged against medical advice.

Strain JNQH373 was isolated from a blood culture of a male patient in his early sixties who had been admitted to a tertiary hospital in Eastern China in August 2019. The patient had a prior medical history of endovascular embolization for an arteriovenous malformation and evacuation of an intracranial hematoma. Upon admission, he presented with fever, respiratory distress, and dyspnea, necessitating orotracheal intubation and mechanical ventilation. Subsequently, a tracheostomy was performed, and closed thoracic drainage was administered. On hospital day 36, the patient experienced a sudden high fever, and blood cultures subsequently recovered a CRKp isolate, JNQH373. Antimicrobial therapy, including tigecycline and imipenem was administered. However, the patient’s health condition continued to deteriorate. On hospital day 39, the patient was discharged at the request of his family for personal reasons.

### Genomic characteristics of the CR-hvKp strains

The genomes of JNQH97 and JNQH373 were sequenced to closure using the combination of Illumina short-reads and Oxford Nanopore long-reads sequencing. Genomic analysis showed that JNQH373 contains one circular chromosome and two plasmids of 55 kb and 280 kb, respectively. In contrast, JNQH97 harbors one chromosome and four plasmids, ranging from 83.3 to 217 kb. In JNQH373, the chromosomal *ybt1* (yersiniabactin siderophore biosynthesis) and *clb* (colibactin toxin biosynthesis) loci were located within ICEKp10. In contrast, JNQH97 harbored the *ybt3* allele in ICEKp3 on its chromosome. JNQH373 belonged to sequence type 23 (ST23) and carried the KL1 capsule and O2a O antigen, while JNQH97 belonged to ST11 and harbored the KL64 capsule type and O1ab O antigen. In JNQH97, the *ompK35* gene was truncated, and a di-amino acid insertion (Glycine-Aspartate) was found inserted in the extracellular loop 3 (L3) region of the OmpK36 protein. In contrast, both *ompK35* and *ompK36* were wild-type in JNQH373.

JNQH373 carried the *bla*_NDM-1_ carbapenemase gene on an IncX3 plasmid (pJNQH373-2), while JNQH97 harbored the *bla*_CTX-M-65_*, bla*_KPC-2_, and *bla*_SHV-12_ gene on an IncR plasmid (pJNQH97-2). *bla*_LAP-2_, *qnrS1*, *sul2*, *tet(A)* were found in another IncFII plasmid (pJNQH97-3) in JNQH97. Both strains contain the pVir plasmids (pJNQH373-1 and pJNQH97-1) with highly conserved plasmid synteny and structure (>99% nucleotide identity, 93% coverage) to the prototype virulence plasmid pK2044. The two pVir plasmids carry the IncHI1B and IncFIBk plasmid replicons, with sizes of 217,079 bp and 279,569 bp, respectively (Table 2). Both pVir plasmids carried the *rmpADC* and *rmpA2D2* loci, however, *rmpA2* genes were truncated in both plasmids (Fig. 1). The aerobactin-encoding *iuc* and *iut* loci were present in both plasmids. However, the salmochelin-encoding *iro* gene cluster was absent in JNQH97. The deletion of the *iroBCDN* gene cluster had been suggested an evolutionary benefit to enhance the survival capacity while maintaining its virulence (13). Additionally, a complete *ter* operon, which confers resistance to tellurite oxide (K₂TeO₃), was identified in both pVir plasmids.

**Fig 1.**
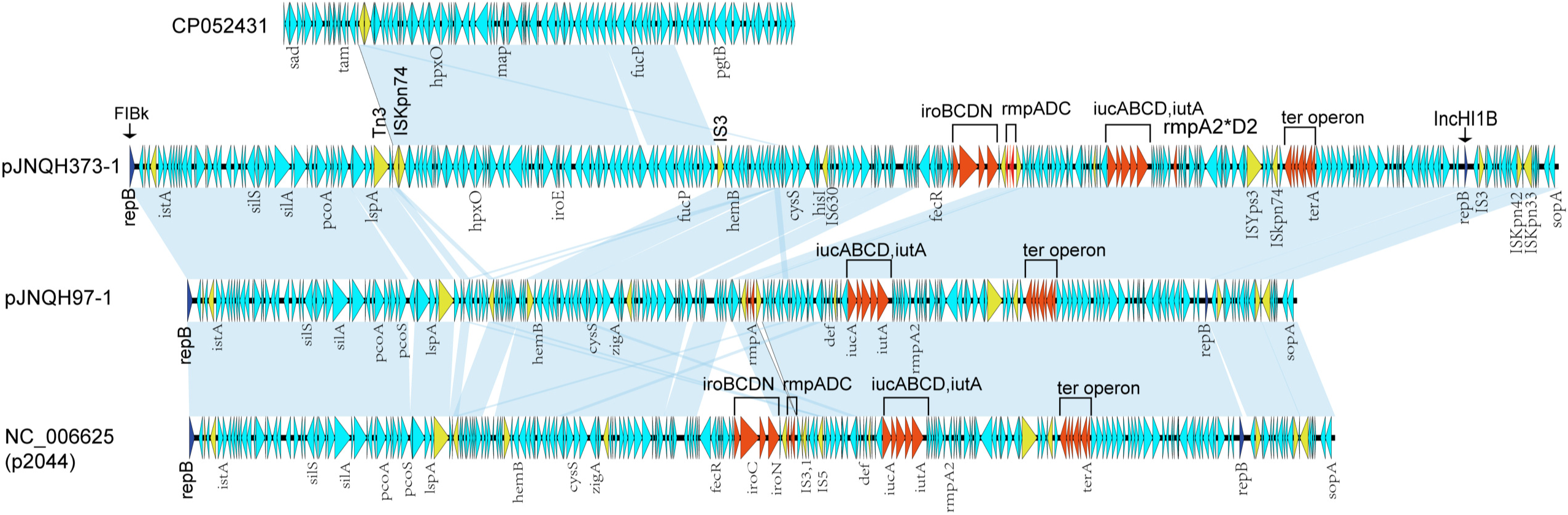
Structural comparison of the pVir-like plasmids from hvKp strains using complete plasmid sequences. *rmpA, rmpA2*, virulence-coding genes (*iro, iuc* and *iut*) and *ter* operons were highlighted in red. Asterisks (*) indicate genes truncated due to premature stop codons. Shallow ocean shading indicates shared regions of homology. ORFs are represented by arrows and are coloured based on predicted gene functions. Light blue arrows, plasmid scaffold regions; dark blue arrows, replication-associated genes; red arrows, virulence-coding genes; and yellow arrows, IS genes. The plasmid accession numbers are listed below the plasmid names.

**Table 2.** Genetic characterization of ST11 and ST23 hvKp strains in this study.

| Strain | Plasmid or chromosome | total_size | Inc type | Resistance and virulence genes |
| --- | --- | --- | --- | --- |
| JNQH97 | Chrome | 5,465,676 |  | *aadA2*, *bla* <sub>SHV-11</sub>, *ybt3*, *ompK35* * |
|  | pJNQH97-1# | 217,079 | IncHI1B/FI Bk | *iucABCD*, *iutA*, *rmpADC*, *rmpA2* *D2 |
|  | pJNQH97-2 | 140,846 | IncR_1/IncF II | *bla*<sub>CTX-M-65</sub>, *bla*<sub>KPC-2</sub>, *bla*<sub>SHV-12</sub> |
|  | pJNQH97-3 | 83,385 | IncFII | *bla*<sub>LAP-2</sub>, *qnrS1*, *sul2*, *tet(A)* |
|  | pJNQH97-4 | 11,970 | ColRNAI |  |
| JNQH373 | Chrome | 5,359,407 |  | *bla*<sub>SHV-11</sub>, *ybt1*, *clb*, *ompK35*, *ompK36* |
|  | pJNQH373-1# | 279,569 | IncHI1B/FI Bk | *iroBCDN*, *iucABCD*, *iutA*, *rmpADC*, *rmpA2* *D2 |
|  | pJNQH373-2 | 55,296 | IncX3_1 | *bla*<sub>NDM-1</sub>, *bla*<sub>SHV-12</sub> |

Further comparative genomics showed these plasmids carried multiple insertion sequences, resulting in extensive rearrangement of pVir genetic elements. Interestingly, pJNQH373-1 contained a 62.1k bp region, flanked by IS*5* family transposase gene IS*Kpn74* and IS*3* family transposase IS*Ec36*, replacing a 10.9k bp fragment. BLASTn analysis indicated this region was nearly identical (100% coverage, 100% identity) to the *K. pneumoniae* chromosome of strain C16KP0122 (accession number CP052431).

However, the fragment was not found in the chromosome of the host JNQH373 strain. The results suggested that the pVir could exchange its elements with the *K. pneumoniae* chromosome and may transfer between different host strains.

### Impact of pVir curing on the in vitro fitness and virulence characteristics of CR-hvKp

To assess the phenotypic and transcriptomic impact of the pVir on the bacterial host, we used our CRISPR-Cas-mediated pCasCure system (27) to precisely eliminate the pVir from the two strains. The complete curing of the pVir was verified by PCR screening using four primer sets targeting specific genes on the pVir plasmids, including *rmpA*, *iucA*, *iroN*, and the HI1B replicon gene (28). Additionally, S1-PFGE analysis verified the successful removal of pVir, demonstrating the absence of the plasmid (Fig. 2A). Lastly, next generation sequencing confirmed the precise curing of pVir in both strains, without additional off-target mutations.

**Fig 2.**
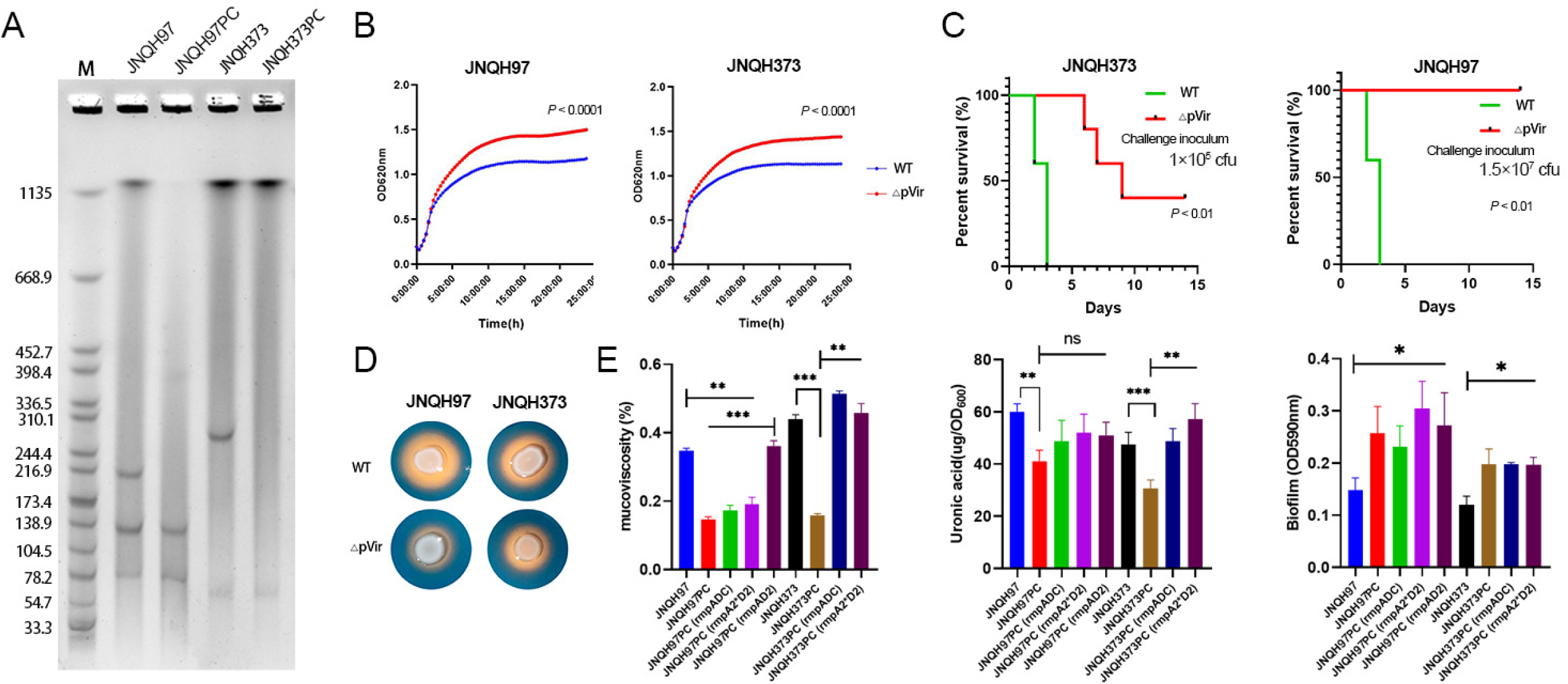
Phenotypic changes of hvKp strain after virulence plasmid curation and introduction of individual rmp locus. (A) S1-PFGE result of the pVir cured and their parental strains. “PC” denotes the pVir cured strains. (B) Growth curve traces for wide-type and pVir cured hvKp isolates. (C) Lethality assay of *Klebsiella pneumoniae* strains in a BALB/c murine infection model. The green line represented WT hvKp strains and the red lines represented their pVir cured strains. The mice were challenged intraperitoneally with challenge inocula of 1×10^5^ CFU for the paired WT/pVir-cured JNQH373, and 1.5×10^7^ CFU for the paired WT/pVir-cured JNQH97 strains. Each group consisted of five mice. The mortality of mice was observed over 14 days. (D) Siderophore secretion experiment on CAS-Agar of hvKp isolates and their pVir cured mutants. Light yellow areas around colonies indicate siderophore secretion. (E) Mucoviscosity assay, uronic acid and biofilm assay of hvkp strains carrying different virulence loci. Each data point was repeated three times (n = 3). ns, not significant, *, *P*<0.05, **, *P*<0.01, ***, *P*<0.001.

We first assessed the *in vitro* fitness costs by comparing the growth curves of pVir-cured and wild-type strains. As shown in Figure 2B, parental strains harboring the pVir plasmids exhibited slower growth rates than their pVir-cured counterparts in both JNQH97 and JNQH373 (*P* < 0.001), suggesting that pVir imposes a significant fitness burden *in vitro*. The string test revealed that parental hvKp strains formed a viscous string of >5 mm, whereas pVir-cured strains lost the HMV phenotype in both JNQH97 and JNQH373. These observations were consistent with the sedimentation assay results, where the pVir-cured mutants of JNQH97 and JNQH373 exhibited over a 50% reduction in mucoviscosity levels in comparison to their respective parental strains, as measured by OD600 (Fig. 2E).

Similarly, both pVir-cured strains displayed reduced siderophore production (Fig. 2D). Uronic acid quantification analysis revealed a ∼35% reduction in uronic acid levels in both JNQH97 and JNQH373, indicating that pVir curing led to decreased capsule production (Fig. 2E). In contrast, biofilm production significantly increased in pVir-cured mutants compared to wild-type strains (Fig. 2E) for both strains. Taken together, pVir curing resulted in reduced HMV, lower levels of CPS secretion, diminished siderophore production, and enhanced biofilm formation in both ST23-KL1 JNQH373 and ST11-KL64 JNQH97 strains.

Scanning electron microscopy (SEM) analysis showed that the capsule in the pVir-cured JNQH373 strain appeared entangled and spread out from the bacterial surface, whereas it was either absent or much shorter in the wild-type parental strain (Fig. 3). Transmission electron microscopy (TEM) imaging of pVir-cured JNQH373 showed surface filaments approximately 0.5–2 µm in length, characteristic of typical type 3 fimbriae (29). However, the filaments were absent in the parental JNQH373 strain. In comparison, SEM and TEM analyses of both the wild-type and pVir-cured JNQH97 strains showed no discernible surface filaments, suggesting a fundamental difference in filamentous structure expression between JNQH373 and JNQH97, regardless of the presence or absence of the pVir plasmid.

**Fig 3.**
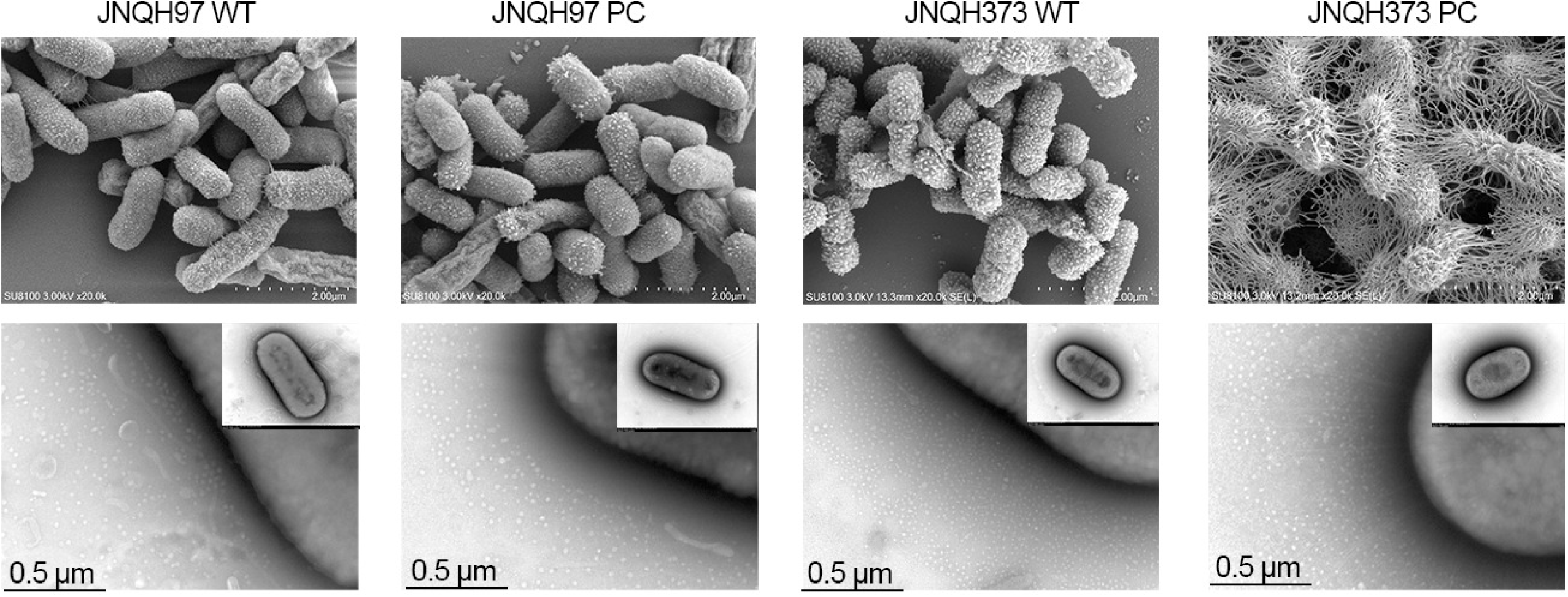
Scanning electron microscopy (SEM) images of wide-type and pVir cured hvKp strains. SEM reveals that a thick variety of filamentous appendages in pVir cured JNQH373. “PC” denotes the pVir cured strains. TEM imaging of JNQH373PC showed surface filaments spread from the surface. However, they were absent in the parental JNQH373 strain. Both the wild-type and pVir-cured JNQH97 strains showed no discernible surface filaments. The scale bar is illustrated at the bottom of the images.

To assess the impact of pVir on *in vivo* virulence, we used a murine lethality assay to evaluate the impact of pVir plasmid curing on the *in vivo* virulence of JNQH373 and JNQH97. Male BALB/c mice were challenged with 1 × 10⁵ CFU of JNQH373 or its pVir-cured mutant and 1.5 × 10⁷ CFU of JNQH97 or its pVir-cured mutant (25, 30) through intraperitoneal injection. At these respective doses, wild-type JNQH373 and JNQH97 caused 100% mortality by day 3. In contrast, pVir-cured JNQH97 did not cause any mortality over a 14-day period, and pVir-cured JNQH373 showed significantly reduced mortality (∼60%, *P*<0.01) compared to wild-type JNQH373 (Fig. 2C). However, the mortality rate of pVir-cured JNQH373 remained significantly higher compared to that of pVir-cured JNQH97, even when the latter was administered at a 150-fold higher dose. This suggests that the unique virulence factors such as the KL (capsular) and O (lipopolysaccharide) antigen types, yersiniabactin in JNQH373, along with other yet unidentified determinants, contribute to its *in vivo* pathogenicity beyond pVir.

### Distinct impact of rmpADC and rmpA2D2 operons on CPS and HMV in CR-hvKp strains

Co-transcription assays confirmed that *rmpADC* and *rmpA2D2* each operate as a single, polycistronic unit under the control of the promoter immediately upstream of *rmpA* or *rmpA2*, respectively. Curing pVir markedly reduced both HMV and CPS production in each background (Fig. 2E). In the ST23-KL1 (JNQH373) derivative, expression of either *rmpADC* or *rmpA2*D2* fully restored mucoviscosity and CPS levels to those of the wild-type. By contrast, in the ST11-KL64 (JNQH97) strain, individual complementation only partially rescued CPS synthesis and failed to recover HMV. Remarkably, introduction of the chimeric *rmpAD2* construct fully restored HMV in ST11-KL64, indicating that RmpA can compensate for the truncated RmpA2, likely a consequence of their close structural similarity (RMSD 2.407 Å, Fig. S1), and that both operons must act together to achieve full hypermucoviscosity in this background. Lastly, all wild-type strains and those complemented with either *rmp* operon exhibited significantly higher biofilm formation compared with their pVir-cured counterparts (Fig. 2E), underscoring the multifaceted role of these regulators in hvKp.

### Impact of pVir curing on the antibiotic susceptibilities of CR-hvKp strains

Susceptibility testing of the pVir-cured and parental strains against several antibiotics was conducted, and the MICs are summarized in Table 3. AST revealed that both parental hvKp strains were resistant to quinolones and most tested β-lactam antibiotics but remained susceptible to amikacin and tobramycin, with MIC values below 2 μg/ml and 1 μg/ml, respectively. JNQH97 was susceptible to ceftazidime/avibactam and aztreonam/avibactam, while JNQH373 showed resistance to ceftazidime/avibactam, consistent with the presence of *bla*_NDM-1_. Additionally, JNQH373 was resistant to tigecycline with an MIC of 4 μg/ml. However, no acquired or mutation mediated tigecycline resistance genes were found. The pVir-cured mutants exhibited MICs similar to their parental strains.

**Table 3.** Minimum inhibitory concentration (MIC) profiles of the pVir cured hvKp and their parental strains (μg/ml).

| Isolates | β-lactams | | | | | | | | | | Aminoglycosides | | Quinolones | | Others | | | |
| --- | --- | --- | --- | --- | --- | --- | --- | --- | --- | --- | --- | --- | --- | --- | --- | --- | --- | --- |
|  | TI<br>M | TZ<br>P | CA<br>Z | SC<br>F | FE<br>P | AT<br>M | IM<br>P | ME<br>M | CA<br>Z-AVI | AT<br>M-AVI | AK | TOB | CI<br>P | LE<br>V | PM<br>B | M<br>H | TG<br>C | SX<br>T |
| JNQH97 | >12<br>8 | >12<br>8 | >64 | >6<br>4 | >3<br>2 | >64 | >1<br>6 | >16 | 4 | 4 | <2 | <1 | >4 | >8 | 2 | >1<br>6 | 2 | <2<br>0 |
| JNQH97P<br>C | >12<br>8 | >12<br>8 | >64 | >6<br>4 | >3<br>2 | >64 | >1<br>6 | >16 | 4 | 4 | <2 | <1 | >4 | >8 | 2 | >1<br>6 | 2 | <2<br>0 |
| JNQH373 | >12<br>8 | >12<br>8 | >64 | >6<br>4 | >3<br>2 | >64 | >1<br>6 | >16 | >32 | 1 | <2 | <1 | >4 | 4 | 1 | >1<br>6 | 4 | <2<br>0 |
| JNQH373<br>PC | >12<br>8 | >12<br>8 | >64 | >6<br>4 | >3<br>2 | >64 | >1<br>6 | >16 | >32 | 1 | <2 | <1 | >4 | 4 | 1 | >1<br>6 | 4 | <2<br>0 |

PC: Virulence plasmid cured strains. TIM, ticarcillin-clavulanate; TZP, piperacillin tazobactam; CAZ, Ceftazidime; SCF, cefoperazone-sulbactam; FEP, cefepime; ATM, aztreonam; IMP, imipenem; MEM, meropenem; CAZ-AVI, ceftazidime-avibactam; ATM-AVI, aztreonam-avibactam; AK, amikacin; TOB, tobramycin; CIP, ciprofloxacin; LEV, levofloxacin; PMB, polymyxin B; MH, minocycline; TGC, tigecycline; SXT, sulfamethoxazole-trimethoprim. All tests were performed in duplicate, and each test included three biological replicates.

We then conducted time-kill assay to assess the kinetics for ceftazidime/avibactam (for JNQH97 pair) and aztreonam/avibactam (for both strain pairs), using a 2× MIC antibiotic concentration. Time-kill curves showed that, in both parental and pVir-cured JNQH97, ceftazidime/avibactam and aztreonam/avibactam achieved an approximately 3∼4-log reduction in bacterial numbers within the first 6 hours, which was sustained through 24 hours. Similarly, aztreonam/avibactam exhibited comparable bacterial killing in both parental and pVir-cured JNQH373, with rapid killing in the first 6 hours and sustained activity through 24 hours (Fig. 4).

**Fig 4.**
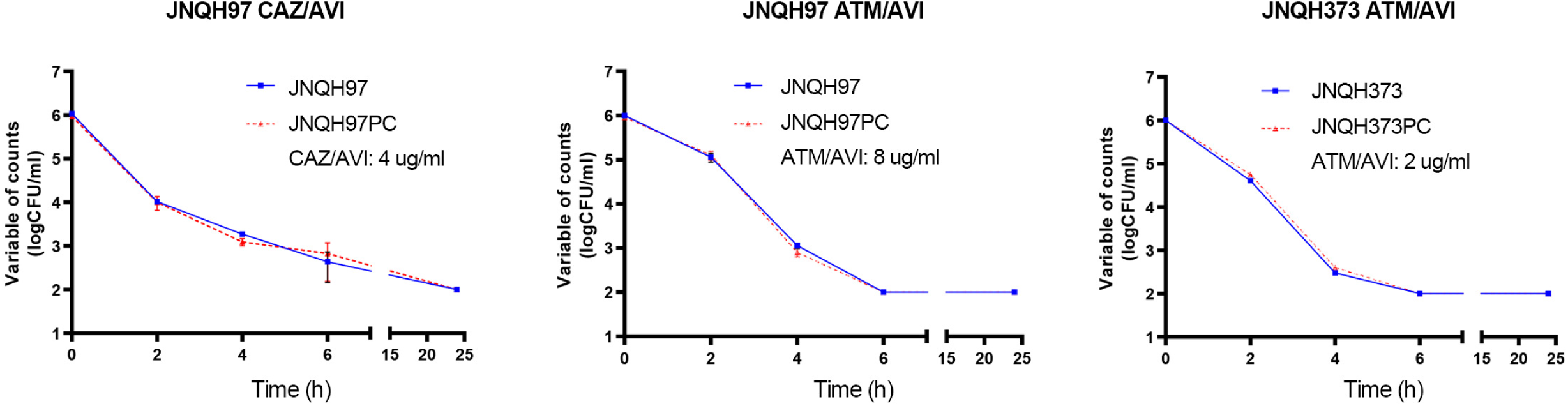
Time-kill curves of parental and pVir -cured hvKp strains exposed to CAZ/AVI and ATM/AVI. “PC” denotes the pVir cured strains. CAZ/AVI and ATM/AVI showed similar bactericidal effects in both parental and pVir-cured strains. Antibiotics concentrations were indicated at each curve. Red dotted lines indicated pVir cured strains while solid blue lines indicated wide-type strains. Data points below the lower limit of detection (100 CFU/mL) were set to 2 log_10_ CFU/mL.

### Impact of pVir curing on genome-wide transcription changes

To assess the transcriptional impact of pVir plasmid curing, RNA sequencing was performed on wild-type and pVir-cured strains of JNQH97 (ST11-KL64) and JNQH373 (ST23-KL1). In JNQH373, 549 differentially expressed genes (DEGs) were identified, comprising 297 upregulated and 252 downregulated genes (Fig. 5A). In contrast, JNQH97 exhibited a more extensive transcriptional response, with 1,182 DEGs—639 upregulated and 543 downregulated. Among the DEGs, 50 were consistently downregulated and 37 upregulated in both strains following pVir curing (Fig. 5B). The shared downregulated genes were primarily involved in CPS biosynthesis, including *galF, manC, wzi*, and genes related to polysaccharide export. These transcriptional changes are consistent with the observed reduction in CPS production and HMV in pVir-cured strains (Fig. 6). However, many DEGs were strain-specific: 471 upregulated and 321 downregulated genes in JNQH97 were not significantly altered in JNQH373, while 188 upregulated and 164 downregulated genes in JNQH373 were not altered in JNQH373. Moreover, 22 genes exhibited opposite expression trends between the two strains.

**Fig 5.**
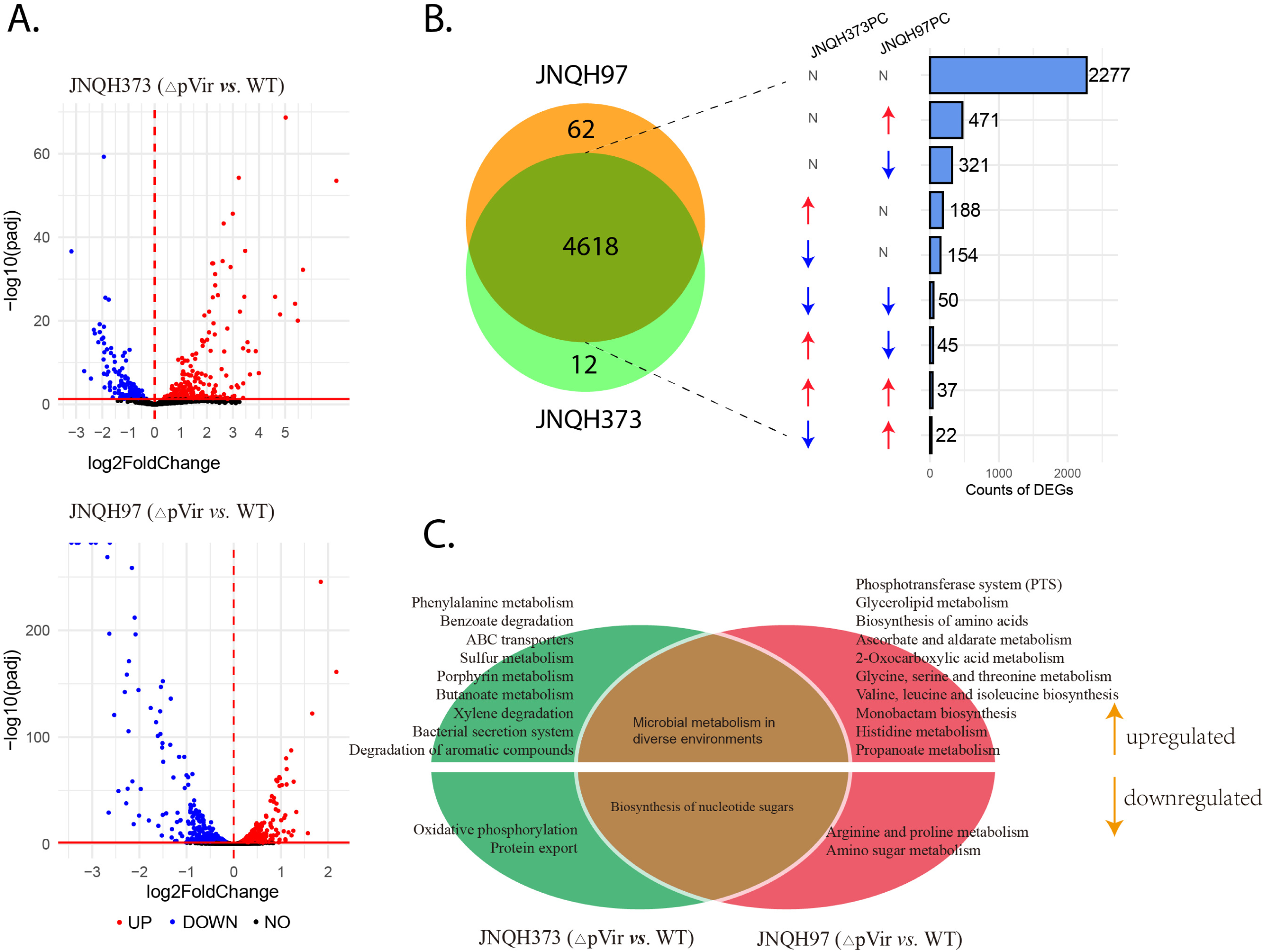
DEGs distribution and enriched metabolism pathways after pVir curing. (A) The volcano plot displays the differential gene expression results from RNA sequencing analysis following pVir curing in hvKp. Upregulated and downregulated genes are shown in red and blue points. (B) Orthologous clusters across JNQH97 and JNQH373 strains. Numbers of strain specific and intersected genes are illustrated. (C) DEGs distribution of pVir-cured mutants relative to wide-type parent strains in JNQH373 and JNQH97 are shown. The numbers of DEGs are labeled on the right of the bar plot. (D) KEGG pathway analysis revealed two pathways commonly affected in both strains following pVir curing. The left oval represents significantly enriched pathways in strain JNQH373, while the right oval represents those in strain JNQH97 following pVir plasmid curing. The overlapping region between the two ovals indicates shared differentially regulated pathways. Pathways located in the upper sections of the ovals are upregulated, whereas those in the lower sections are downregulated.

**Fig 6.**
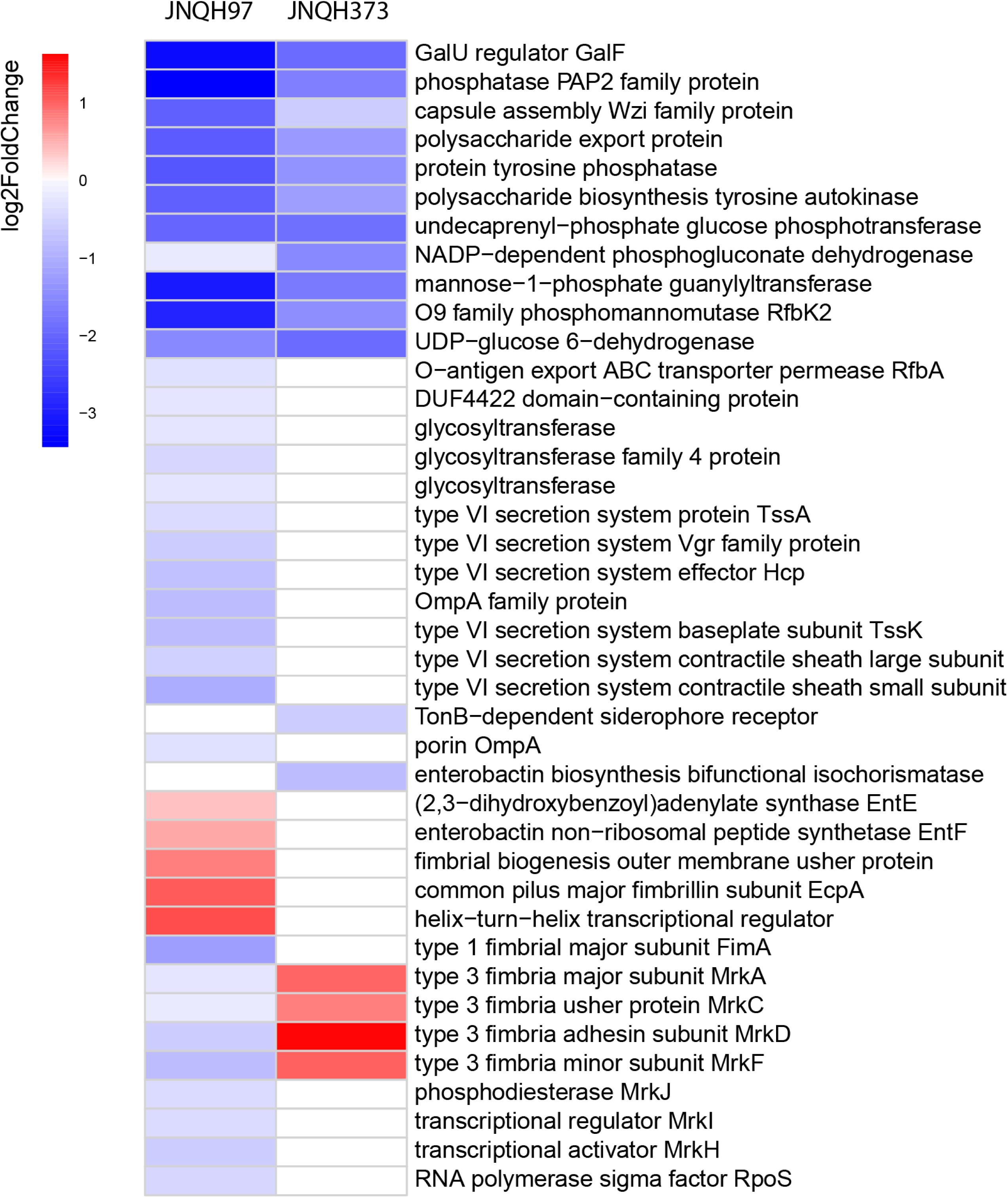
Expression alteration of virulence associated genes after pVir curation. Virulence genes those did not significantly changed were designated as white blocks.

Distinct regulatory patterns were observed for genes related to adhesion and secretion systems. In JNQH373, type III fimbrial genes (*mrkA, mrkC, mrkD*, and *mrkF*) were significantly upregulated, correlating with enhanced biofilm formation and electron microscopy observations. The upregulation probably mediated by the lose of *mrkABCDF* repressor in pVir, iroP (31) or rmpA (32). However, these genes were downregulated in JNQH97, despite increased biofilm production (Fig. 6). This discrepancy may be explained by the upregulation of alternative biofilm-related genes in JNQH97, such as *ecpA* (33) and genes encoding chaperone-usher pathway proteins (34), indicating different mechanisms driving biofilm enhancement across strains. Additionally, genes associated with the type VI secretion system (T6SS) were significantly downregulated in pVir-cured JNQH97 but remained unchanged in JNQH373, further highlighting the strain-specific regulatory responses to pVir loss (Fig. 6).

KEGG pathway analysis revealed two pathways commonly affected in both strains following pVir curing (Fig. 5C). One was upregulated microbial metabolism in diverse environments; and the other was downregulated biosynthesis of nucleotide sugars. Nucleotide sugars are key precursors for the synthesis of bacterial capsular polysaccharides, lipopolysaccharides (LPS), and exopolysaccharides (35), which supports the observed reduction in capsule production.

In addition to these shared changes, strain-specific pathway alterations were identified (Fig. 5C). In JNQH373, pVir curing resulted in the upregulation of several metabolic pathways, including those involved in phenylalanine, sulfur, porphyrin, and butanoate metabolism, while pathways related to oxidative phosphorylation and the protein export system were downregulated. In contrast, in JNQH97, most of the affected pathways were related to amino acid metabolism. Specifically, histidine, glycine, serine, and threonine metabolism were upregulated, whereas arginine and proline metabolism were downregulated. Together, these results highlight both shared and strain-specific transcriptional responses to pVir loss, emphasizing the distinct regulatory landscapes of ST11 and ST23 hypervirulent *K. pneumoniae*.

## DISCUSSION

The emergence of convergent CR-hvKp poses a significant threat to global public health. This study provides critical insights into the role of the virulence plasmid in shaping the phenotypic and transcriptional profiles of two distinct CR-hvKp strains, JNQH373 (ST23-KL1) and JNQH97 (ST11-KL64). By employing a CRISPR-Cas9-mediated pCasCure system (36) to generate isogenic pVir-cured mutants, we elucidated the impact of pVir on bacterial fitness, virulence, and antimicrobial resistance, while also uncovering strain-specific regulatory mechanisms.

Our findings underscore the central role of pVir in maintaining the hypervirulent phenotype of CR-hvKp. The loss of pVir resulted in a significant reduction in HMV, capsule production, and siderophore activity in both JNQH373 and JNQH97, consistent with previous reports that pVir-encoded genes, such as *rmpADC*, *iucABCD*, and *iroBCDN*, are critical for these phenotypes (14, 22). Notably, the pVir-cured strains exhibited reduced virulence in a murine sepsis model, exhibiting significantly lower mortality compared to their wild-type counterparts. These results highlight the indispensable role of pVir in hvKp pathogenesis. However, the remaining virulence observed in pVir-cured JNQH373 suggests that additional chromosomal factors, such as the yersiniabactin and colibactin loci, or different capsule types, contribute to its pathogenicity, particularly in the ST23-K1 lineage.

Our complementation experiments revealed distinct, lineage-specific regulatory roles of the *rmpADC* and *rmpA2D2\** operons in the production of capsular CPS and HMV between ST23-K1 and ST11-KL64 strains. In the ST23-K1 strain JNQH373, either *rmpADC* or *rmpA2D2\** alone was sufficient to restore CPS and HMV levels, indicating functional redundancy between these operons. In contrast, in the ST11-KL64 strain JNQH97, neither *rmpADC* nor *rmpA2D2\** complementation could fully restore CPS or HMV, although *rmpADC* partially increased CPS production. Interestingly, in JNQH97, the *rmpA* gene appears capable of compensating for the truncated *rmpA2* gene, thereby contributing to the HMV phenotype. This compensatory effect may be due to the structural homology between RmpA and RmpA2 proteins. The inability of individual operons to fully restore CPS and HMV in this strain highlights the complexity of capsule regulation in convergent CR-hvKp lineages. These findings support the notion that the regulatory mechanisms governing CPS and HMV expression are lineage-dependent, consistent with recent studies (22).

The presence of the pVir plasmid did not significantly impact the antimicrobial susceptibility profiles of *K. pneumoniae* strains JNQH373 and JNQH97, as both wild-type and pVir-cured variants exhibited comparable minimum inhibitory concentrations (MICs) across most tested antibiotics. Notably, time-kill assays demonstrated that ceftazidime/avibactam maintained similar bactericidal activity against both wild-type and pVir-cured JNQH97 strains, which harbor the *bla*_KPC_ resistance gene. Likewise, aztreonam/avibactam showed equivalent efficacy against wild-type and pVir-cured JNQH373 strains carrying the *bla*_NDM_ gene. These results suggest that ceftazidime/avibactam and aztreonam/avibactam may serve as effective therapeutic options for treating infections caused by susceptible hypervirulent *K. pneumoniae* strains.

RNA sequencing revealed extensive transcriptional changes following pVir curing, with JNQH97 exhibiting a greater number of DEGs than JNQH373. This finding suggests that pVir imposes a more significant regulatory burden on the ST11-KL64 background, likely reflecting its more recent acquisition of pVir compared to the ST23-KL1 lineage. The downregulation of CPS biosynthesis genes in both strains correlated with the observed reduction in capsule production and HMV reduction, while the upregulation of biofilm-associated genes, such as type III fimbriae in JNQH373 and chaperone/usher pathway proteins in JNQH97, aligned with the increased biofilm formation in pVir-cured strains. Notably, genes linked to the type VI secretion system were markedly downregulated in JNQH97 following pVir curing, whereas their expression levels remained unchanged in JNQH373. These strain-specific transcriptional responses highlight the adaptability of *K. pneumoniae* to pVir loss and underscore the diverse regulatory mechanisms governing virulence and persistence.

The convergence of hypervirulence and multidrug resistance in *K. pneumoniae* represents a formidable challenge for clinical management. Our findings emphasize the need for targeted therapeutic strategies that address both virulence and resistance mechanisms. For instance, the development of inhibitors targeting pVir-encoded virulence factors, such as aerobactin biosynthesis or the *rmpA* regulated capsule synthesis network, could attenuate hvKp pathogenicity without exerting selective pressure for resistance. Additionally, the strain-specific differences in CPS regulation warrant further investigation to identify lineage-specific vulnerabilities that could be exploited for treatment.

A limitation of this study is the inclusion of only two CR-hvKp strains representing ST23 and ST11 lineages. While these sequence types are clinically relevant and genetically distinct, the findings may not fully capture the diversity of virulence plasmid–host interactions across the broader spectrum of hvKp strains with different genetic backgrounds. Further research involving a wider range of sequence types is needed to determine whether virulence plasmids exert consistent or strain-specific effects on bacterial physiology and pathogenicity.

In conclusion, this study provides a comprehensive understanding of the role of pVir in CR-hvKp strains, highlighting its impact on virulence, fitness, and antimicrobial resistance. The strain-specific regulatory networks uncovered here underscore the complexity of *K. pneumoniae* pathogenesis and the need for tailored approaches to combat this evolving pathogen. Future research should focus on elucidating the molecular mechanisms underlying pVir-mediated virulence and resistance, as well as exploring novel therapeutic interventions to mitigate the threat posed by CR-hvKp.

## MATERIALS AND METHODS

### Ethical statement

This study was conducted in accordance with the principles of the Declaration of Helsinki. Ethics committee approval of this study was obtained from the institutional review board (IRB) of the First Affiliated Hospital of Shandong First Medical University [ethics approval number YXLL-KY-2023(025)]. This project did not affect the normal diagnosis and treatment of the patient; after consultation with the IRB of the First Affiliated Hospital of Shandong First Medical University, formal written patient consent was not required. For animal-related experiments, the animal protocol was reviewed and approved by the Animal Ethics Committee of the First Affiliated Hospital of Shandong First Medical University (animal ethics approval number: S078).

### DNA sequencing and bioinformatics

Genomic DNA was isolated using a Wizard® HMW DNA Extraction Kit (Promega, Madison, WI, United States) and was subjected to Illumina Nextseq and Nanopore MinION sequencing. Hybrid de novo assembly was conducted using Unicycler v0.49 (37). The whole-genome sequences were annotated by Prokka v 1.14.6 (38) followed by manual curation. Kleborate v. 2.2 (39) was used for *Klebsiella* multi-locus sequence typing, K locus and O locus typing. The antibiotic resistance/plasmid replicon genes were identified out using ABRicate v.1.0.1 (https://github.com/tseemann/abricate) using CARD (40) and PlasmidFinder (41) database, respectively. Virulence locus (yersiniabactin, aerobactin, and other siderophore production systems) were identified using Kleborate v2.2 in combination with VFDB database (42). The complete sequences of virulence plasmids were compared with plasmids pK2044 (NC_006625.1), from K1 prototype strain NTUH-K2044 (43), using BLASTn and illustrated by Easyfig (44).

### CRISPR/Cas9-mediated pVir plasmid curing

CRISPR/Cas9-mediated pVir plasmid curing was conducted following our previously described method with minor modifications (36). Briefly, the 20 nt base-pairing region of an sgRNA targeted IncHI1B plasmid replicon was designed using the R package CRISPRseek (45). The sgRNA was introduced into pCasCure backbone using PCR, restriction enzyme digestion, and ligation, as previously described (resulting in the pCasCure-N20_HI1B_ plasmid). The pCasCure-N20_HI1B_ was electroporated into the hvKp strain, followed by 1% arabinose treatment for 6h at 37℃ with shaking (200rpm). The culture was plated on LB agar containing apramycin and then checked for the loss of targeted pVir plasmid by PCR. The pVir cured strains were then streaked onto a LB agar plate containing 5% sucrose to remove pCasCure-N20_HI1B_.

### S1 pulsed-field gel electrophoresis (PFGE)

To test the clearance of the pVir plasmid through the pCasCure system, we used S1 PFGE to confirm the successful curing of pVir. The parental and pVir cured hvKp strains were digested with S1 nuclease after being embedded in 1% SeaKem Gold Agarose gels, followed by plasmid DNA separation by PFGE as described previously (46). *Salmonella enterica* serotype Braenderup H9812 was used as a size marker. pVir cured strains were also subject to Illumina NextSeq sequencing as described above to check potential mutations through plasmid curing.

### Co-transcription analysis

To assess whether the *rmpACD* and *rmpA2D2* operons were respectively co-transcribed as single RNA, total RNA was isolated from strains JNQH97 and JNQH373 using the Bacteria RNA Isolation Kit (YALEPIC, China), following the manufacturer’s instructions. Subsequently, 200 ng of DNase-treated RNA was reverse transcribed into complementary DNA (cDNA) using the All-in-Mix 1^st^ Strand cDNA Synthesis Kit (YALEPIC, China). Primer sets targeting the junctions of *rmp* operon components were used to perform PCR amplification on cDNA, genomic DNA (gDNA), and DNase-treated RNA controls. These reactions served to confirm both operon co-transcription and the absence of gDNA contamination. Promoter regions upstream of the *rmpA* and *rmpA2* genes were predicted using the BPROM tool (47).

### Cloning of plasmid-borne rmp homologues

The *rmpA*, *rmpA2* loci and the upstream sequences were amplified with primer pairs rmpADC_locF/R and rmpA2D2_locF/R respectively, using plasmid DNA of strains JNQH373 and JNQH97 as the templates, respectively. These amplified fragments were ligated to *pEASY*^®^-T1 Cloning Vector (TransGen, China) by TA cloning and then transformed into *E. coli* strain DH5α competent cells. The resulting plasmids recovered from the transformants were verified by Sanger sequencing and then transformed into pVir cured JNQH97 and JNQH373, respectively. To examine the combined effect of *rmpA* and *rmpD2* in strain JNQH97, a chimeric *rmpAD2* construct was generated by PCR amplifying the two genes separately with primer pairs rmpADC_locF/rmpA_homo and rmpA2_homo/rmpA2_locR (Fig. S2). The two fragments were assembled by seamless clone then introduced into the pVir-cured recipient cells. The primers used for cloning are listed in Table 4.

**Table 4.**
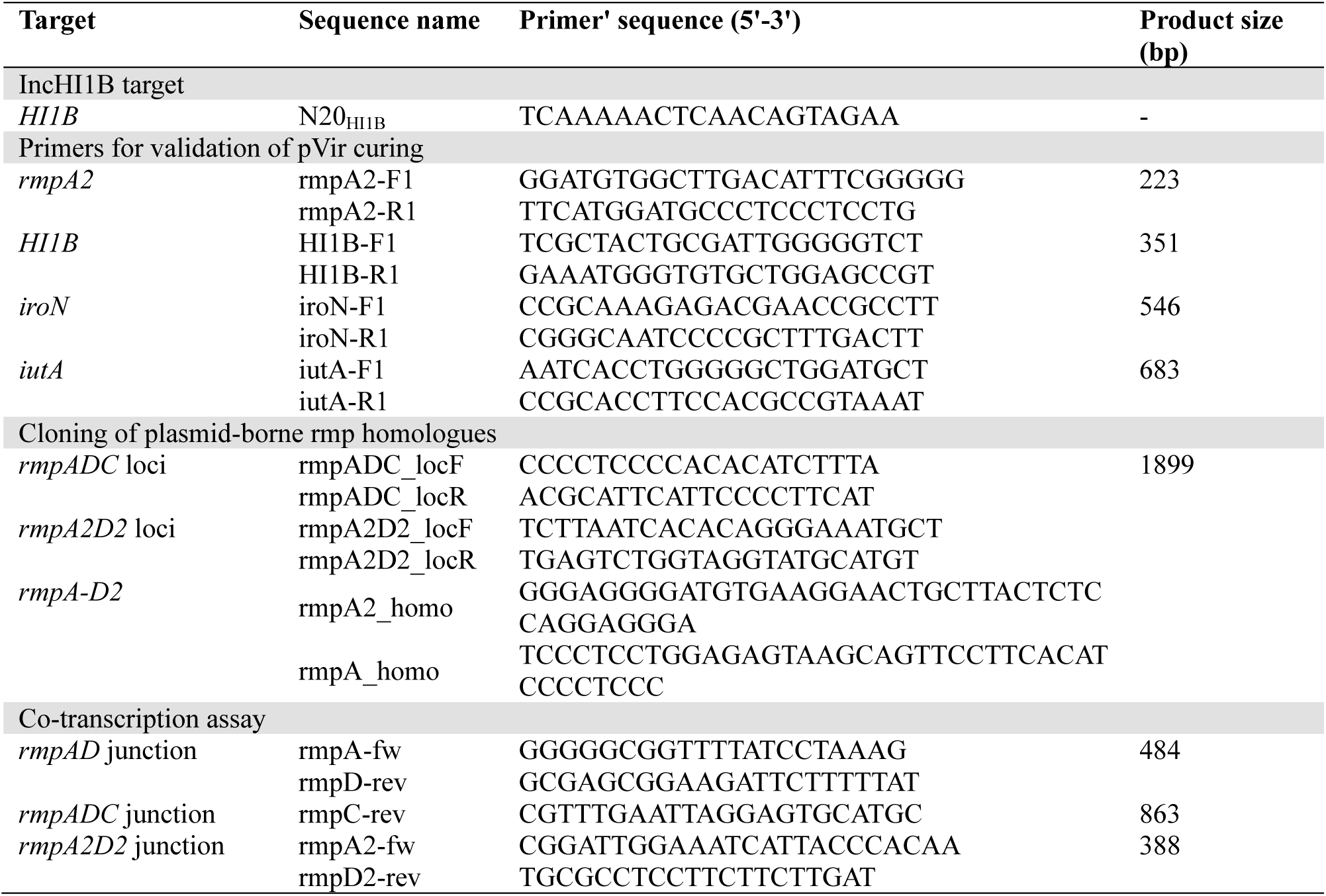
Primers and plasmids used in this study.

### Structural similarity assay

To evaluate structural similarity between the RmpA and RmpA2 proteins, 3D models were generated using AlphaFold (48), based on the wild-type amino acid sequences. Structural alignment of the predicted models was performed using PyMOL (49), and the root mean square deviation (RMSD) value was calculated to quantify similarity between the aligned protein structures (50).

### Hypermucoviscosity assay

The mucoviscosity of the strains was determined using a string test and a sedimentation assay. A string test was performed by stretching the bacterial colonies on sheep blood agar plates using an inoculation loop. Strings of 5 mm or longer were defined positive. For the sedimentation assay, cultures of *K. pneumoniae* were grown overnight in 5 ml LB medium at 37℃. The optical density at 600 nm (OD600) of cultures were measured, followed by spinning down at 2,500g for 5 min. OD600 of the top 1 ml of supernatant was measured, and the sedimentation results were expressed as a ratio of the supernatant OD600 to that in the input culture.

### Growth Curves

To evaluate the fitness impact of the pVir plasmid, bacterial growth was monitored in Mueller-Hinton (MH) broth. Overnight cultures were pelleted and washed twice with PBS, then resuspended and adjusted to an initial optical density at 600 nm (OD_600_) of 0.005 in fresh MH medium. Aliquots of 200 μL from each prepared isolate were dispensed into individual wells of a sterile 96-well microtiter plate. Growth was tracked over a 24-hour period at 37 ℃ using a Multiskan™ FC Microplate Photometer (Thermo Scientific, IE), with OD_600_ readings recorded at regular intervals. Each strain was tested in triplicate per experiment, and the entire procedure was independently repeated three times to ensure reproducibility.

### Antibiotic susceptibility testing (AST)

AST was performed using the automated VITEK 2 system (bioMérieux, Nürtingen, Germany) as described in our prior work (51). For ceftazidime-avibactam and aztreonam-avibactam, MICs were assessed via broth microdilution, with avibactam maintained at a fixed concentration of 4 μg/mL. MIC values were interpreted in accordance with the Clinical and Laboratory Standards Institute (CLSI) guidelines (M100-S32). Quality control was ensured using reference strains *E. coli* ATCC 25922 and *P. aeruginosa* ATCC 27853. Each assay was conducted in triplicate across two independent days to confirm reproducibility. CLSI breakpoints were applied for all agents except tigecycline, for which the EUCAST epidemiological cutoff of >2 μg/mL was used.

### Biofilm formation assays

Biofilm formation was measured by crystal violet staining. In brief, overnight cultures were diluted 1:1,000 in LB medium and a total volume of 200μl was transferred to a vinyl “U”-bottom 96-well microtiter plates. Following incubation at 37 ℃ for 24h, the wells were washed four times with distilled water to rinse away the planktonic bacteria. The remaining adherent bacteria are stained with 125 μl of 0.1% crystal violet dye. After 10 min incubation, crystal violet was removed and the wells were washed six times with distilled water. the stained biofilms are solubilized 150 μl 30 % glacial acetic acid in water and the plate was incubated for 10 min at room temperature before the OD590 measurement with a microplate reader (Thermo Scientific, IE). At least three replicates were used for each sample, and the experiments were repeated twice.

### Siderophore secretion

Strains for their ability to secrete siderophores were analyzed using the blue agar chrome azurol S (CAS) assay (52). Briefly, overnight cultures were diluted 1:100 in 5 ml fresh LB medium and grown to an OD600 of 0.6. 5 µl cultures were then put on agar plates containing chrome azurol S-iron(III)-hexadecyltrimethylammonium bromide and incubated overnight at 37 ℃. Iron uptake was determined visually by color change from blue to yellow on the following day.

### Capsule isolation and quantification

Overnight bacterial cultures (500 μL) were combined with 100 μL of 1% Zwittergent 3-14 (Aladdin) prepared in 100 mM citric acid buffer (pH 2.0). The mixture was incubated in a shaking water bath at 50 ℃ and 170 rpm for 30 minutes. Following incubation, samples were centrifuged at 16,000 × g for 2 minutes. 300 μL supernatant was transferred to fresh 1.5 mL microcentrifuge tubes, and CPS was precipitated by adding 1.2 mL of absolute ethanol. Tubes were placed on ice for 30 minutes, then centrifuged at 16,100 × g for 10 minutes at 4 ℃. Pellets were air-dried for 30 minutes and resuspended in 100 μL of distilled water, followed by overnight incubation at room temperature.

Quantification of CPS was performed by measuring uronic acid content. For each sample, 20 μL of resuspended CPS was mixed with 120 μL of 12.5 mM sodium tetraborate (Aladdin) in concentrated sulfuric acid in PCR tubes. A standard curve was generated using serial dilutions of galacturonic acid (0– 100 μg/mL, Aladdin). Tubes were heated at 100 ℃ for 5 minutes with intermittent shaking, then cooled at room temperature for 15 minutes. Subsequently, 2 μL of 0.15% 3-phenylphenol (Aladdin) in 0.5% NaOH was added, and samples were allowed to equilibrate for another 15 minutes. After a final 5-minute incubation with shaking, reactions were transferred to a 96-well microplate. Absorbance was recorded at 520 nm using a Multiskan™ FC Microplate Photometer (Thermo Scientific, IE). CPS concentrations were extrapolated from a glucuronic acid standard curve. All assays were performed in triplicate to ensure reproducibility.

### Time-kill experiments

Time-kill experiments were performed at 2× MIC with CAZ/AVI (4 μg/ml), ATM/AVI (8 μg/ml for JNQH97, 2 μg/ml for JNQH373) for the susceptible strains. For CAZ/AVI and ATM/AVI testing, avibactam concentration was fixed at 4 mg/L. Precultures were prepared to achieve starting inocula of 10^6^ CFU/mL in a total volume of 1.5 mL. The tubes were incubated on a shaker (200 rpm) at 37℃. Samples for viable counts were taken at 0 (before the addition of antibiotics), 2, 4, 6 and 24h. 10μL samples were serially diluted as appropriate and plated onto Mueller-Hinton agar plates, which were placed in an incubator at 37℃ for 24 h. Data points below the lower limit of detection (100 CFU/mL) were set to 2 log_10_ CFU/mL.

### In vivo murine lethality assay

The procedure for the murine lethality assay followed a previously described method (53). Male BALB/c mice, aged 6 to 8 weeks, were purchased from Beijing Vital River Laboratory Animal Technology Co., Ltd. (Beijing, China) and challenged intraperitoneally with a 1-mL sterile syringe. The first inoculation consisted of 1×10^5^ CFU in 100 μl of 1×PBS for all wild-type or pVir-cured hvkp strains. Since no lethality was observed in the WT/pVir-cured strain JNQH97, the challenge inoculum was increased to 1.5×10^7^ CFU. The survival rate of the mice was observed daily for 14 days, and GraphPad Prism 8 was used to generate survival curves. The log-rank (Mantel–Cox) test was used for statistical analysis. Each experimental group consisted of five mice. To minimize pain and distress, mice were monitored at least twice daily following intraperitoneal bacterial inoculation and were humanely euthanized by cervical dislocation if they exhibited signs of severe illness or distress such as lethargy, ruffled fur, or labored breathing.

### RNA sequencing (RNAseq) transcriptome analysis

RNAseq was conducted in pVir cured and their parent strains to investigate the impact of pVir plasmid on hvKp bacterial host. Three replicates of each strain were performed. The bacterial strains were grown to mid-log phase at 37℃ in LB broth. Cells were harvested and treated with RNAprotect (Qiagen), followed by extraction using UltraClean RNA isolation kits (MoBio). cDNA libraries were constructed with ScriptSeq Complete Gold kits (Epicentre Biosciences) and were sequenced on an Illumina HiSeq instrument. Raw reads from the sequenced libraries were subjected to quality control (QC) to trim the adaptor sequences using Trim Galore (version 0.4.5), and to filter out RNA sequences with sormerna (54). The clean reads were mapped to the complete JNQH373 and JNQH97 genomes using Rsubread package (55). FeatureCounts function within the Rsubread package was used to summarize the data to gene-level read counts. Transcript abundance data were conducted using the software package DESeq2 (56). The shrunken log fold changes was estimated using the method of approximate posterior estimation for generalized linear model (apeglm) (57). Differentially expressed genes were identified using an adjusted *P* value (padj) threshold of < 0.05. Kyoto Encyclopedia of Genes and Genomes (KEGG) pathway annotation of differentially expressed genes were achieved by KOBAS-intelligence (58). Gene expression levels were subject to Gene Set Enrichment Analysis (GSEA) analyses using the clusterProfiler 4.0 (59). Comparative analysis of orthologous clusters across JNQH97 and JNQH373 strains were determined by OrthoVenn2 web server (60).

### Scanning and transmission electron microscopy

For scanning microscopy, overnight cultures of bacteria were harvested at 2,000 × g for 7 min and washed with 1× PBS (pH 7.3). The bacteria were then fixed in 2.5% glutaraldehyde (Sigma-Aldrich) in 1× PBS (pH 7.3) for 2 h at 4℃. Then the samples were rinsed in 1× PBS and subsequently post-fixed in the solution of osmium tetroxide at 4℃. After washing in distilled water, the samples were dehydrated in a series of ethanol of ascending concentrations up to 100 % and dried at the critical point of liquid CO_2_. Lastly, the samples were mounted on aluminum specimen stubs using carbon adhesive tapes, sputter-coated with a 15 nm gold/palladium layer and examined using scanning electron microscopy. For transmission electron microscopy, the bacteria were deposited onto a copper grid coated with a carbon film and allowed to adhere for 3 to 5 minutes. Subsequently, they were subjected to staining using a 2% phosphotungstic acid solution for 1 to 2 minutes. Following this, the copper grids were air-dried at room temperature before being examined using transmission electron microscopy.

### Statistical analysis

An unpaired two-sided Student’s *t*-test was performed to analyze the statistical difference between the levels of viscosity, biofilm formation and CPS production. To compare the growth curves between the wild-type and pVir-cured hvKp strains, nonlinear regression analysis was performed using a logistic growth model. The difference in BALB/c murine survival between groups was assessed using the Log-rank (Mantel–Cox) test, which evaluates statistical significance across Kaplan–Meier survival curves. A threshold of *P* < 0.05 was considered statistically significant. The statistics analysis was performed with GraphPad Prism 8.

## Data Availability

The genome sequences of hvKp strains JNQH97 and JNQH373, along with the RNA-Seq data for the pVir-cured mutants and their respective wild-type parental strains, have been deposited in Figshare. For strain JNQH97, raw Illumina RNA-Seq data are labeled as follows: Wild-type: J97_1, J97_2, J97_3; pVir-cured mutant: J97_4, J97_5, J97_6. For strain JNQH373, raw Illumina RNA-Seq data are labeled as: Wild-type: W373_1, W373_2, W373_3; pVir-cured mutant: M373_1, M373_2, M373_3. Each condition includes three biological replicates. The dataset is available at https://doi.org/10.6084/m9.figshare.30446609) (61).

---

**Fig S1.**
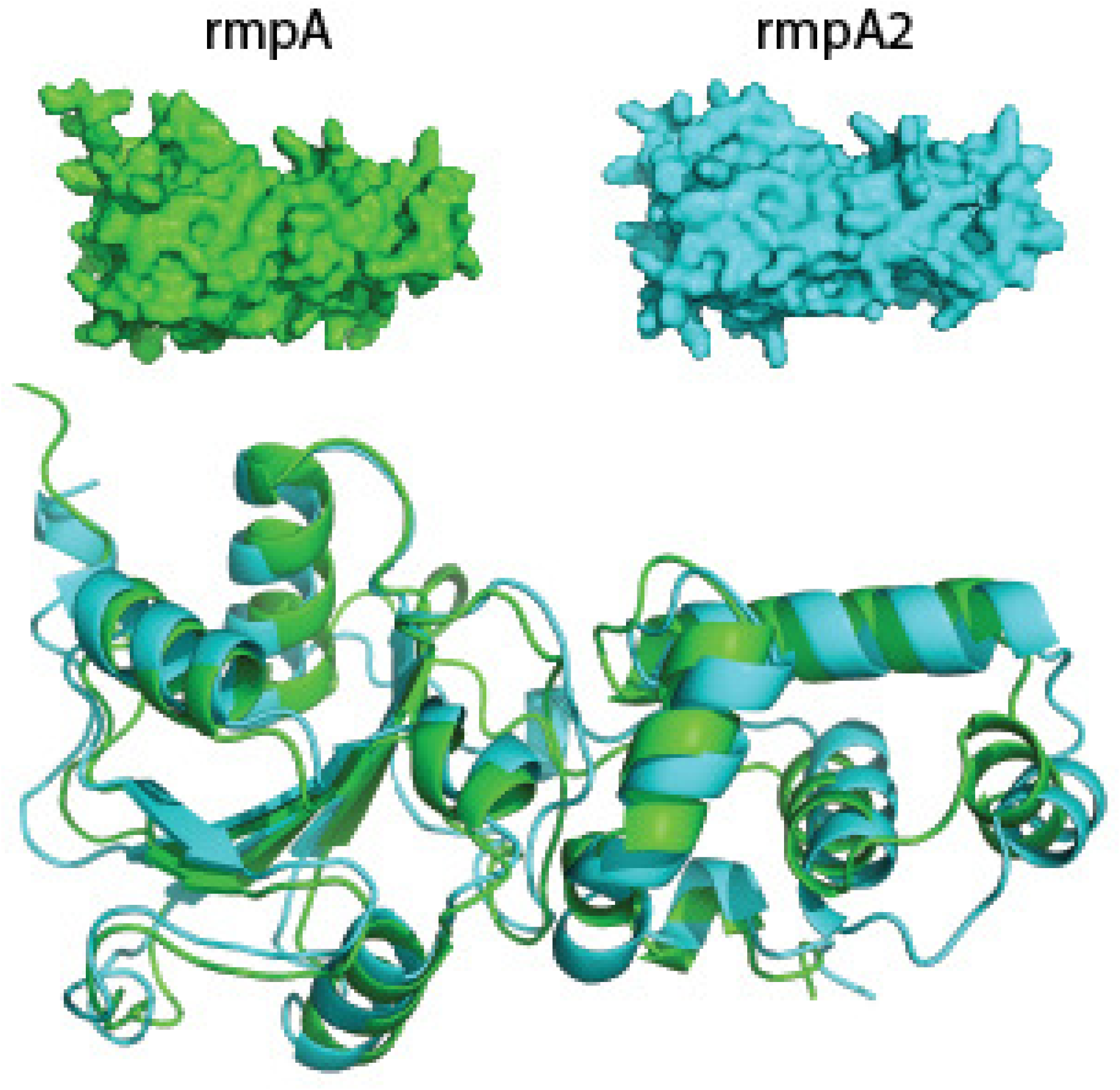
3D structure and similarity alignment of rmpA and rmpA2 protein. Similarity alignment analysis showed rmpA and rmpA2 had close structural similarity with RMSD value of 2.407 Å.

**Fig S2.**
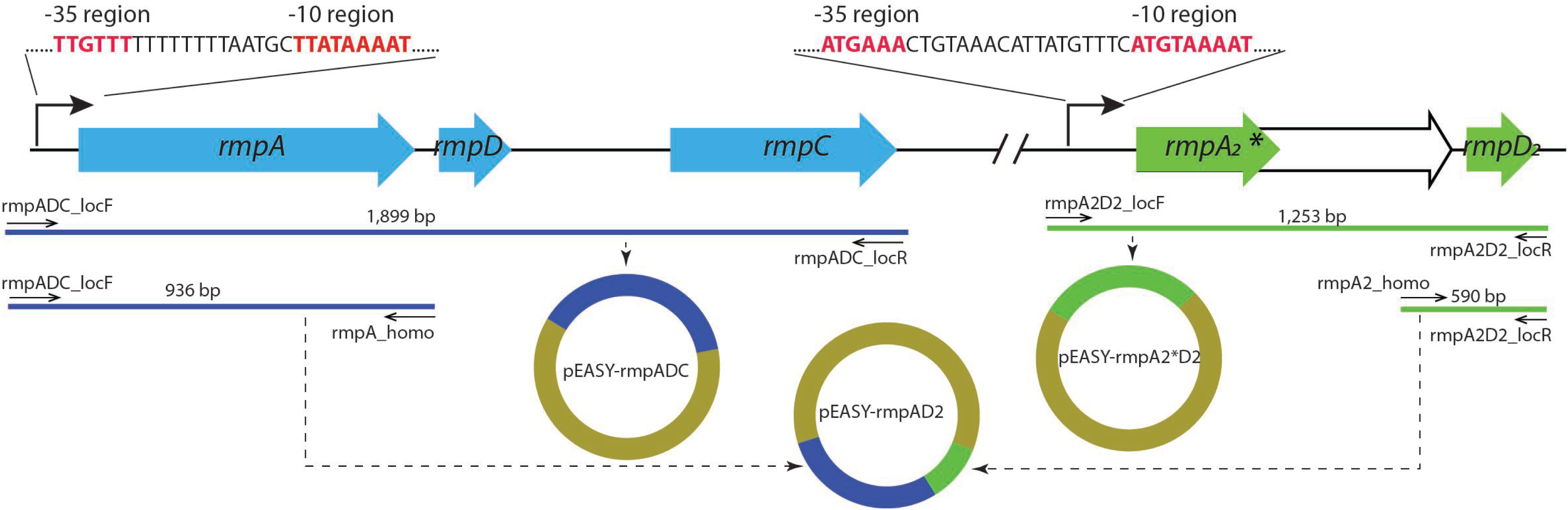
Cloning strategy for *rmpADC*, *rmpA2*D2* and *rmpAD2* operons. Predicted promoter sequences are shown, with the –35 and –10 consensus elements highlighted in red. Primers and the sizes of the PCR fragments are illustrated.

